# Prime Editing in Mice Reveals the Essentiality of a Single Base in Driving Tissue-Specific Gene Expression

**DOI:** 10.1101/2020.11.07.372748

**Authors:** Pan Gao, Qing Lyu, Amr R. Ghanam, Cicera R. Lazzarotto, Gregory A. Newby, Wei Zhang, Mihyun Choi, Orazio J. Slivano, Kevin Holden, John A. Walker, Anastasia P. Kadina, Rob J. Munroe, Christian M. Abratte, John C. Schimenti, David R. Liu, Shengdar Q. Tsai, Xiaochun Long, Joseph M. Miano

## Abstract

Most single nucleotide variants (SNVs) occur in noncoding sequence where millions of transcription factor binding sites (TFBS) reside. Several genome editing platforms have emerged to evaluate the functionality of TFBS in animals. Here, a comparative analysis of CRISPR-mediated homology-directed repair (HDR) versus the recently reported prime editing 2 (PE2) system was carried out in mice to demonstrate the essentiality of a single TFBS, called a CArG box, in the promoter region of the *Tspan2* gene. HDR-mediated substitution of three base pairs in the *Tspan2* CArG box resulted in 20/37 (54%) founder mice testing positive for the correct edit. Mice homozygous for this edit showed near loss of *Tspan2* expression in aorta and bladder with no change in heart or brain. Using the same protospacer, PE2-mediated editing of a single base in the *Tspan2* CArG box yielded 12/47 (26%) founder mice testing positive for the correct edit. This single base substitution resulted in ∼90% loss of *Tspan2* expression in aorta and bladder with no change in heart or brain. Targeted sequencing demonstrated all PE2 and HDR founders with some frequency of on-target editing. However, whereas no spurious on-target indels were detected in any of the PE2 founders, many HDR founders showed variable levels of on-target indels. Further, off-target analysis by targeted sequencing revealed mutations in 5/11 (45%) HDR founders but none in PE2 founders. These results demonstrate high fidelity editing of a TFBS with PE2 and suggest a new paradigm for Cre/*lox*P-free tissue-specific gene inactivation via single base substitution in a TFBS. The PE2 platform of genome editing represents a powerful approach for modeling and correcting relevant noncoding SNVs in the mouse.

---

Proper spatiotemporal control of gene expression requires the RNA polymerase II complex to physically associate with DNA-binding transcription factors and their coregulators over transcription factor binding sites (TFBSs) located in the promoter and enhancer region of target genes.^1^ Elucidating enhancer function and the role of individual TFBSs has informed our understanding of basic mechanisms of gene transcription as well as the development of Cre/*lox*P mouse models for cell-restricted gene inactivation. Further, since most sequence variants (*e*.*g*., single nucleotide variants or SNVs) associated with human disease occur in noncoding sequence space where TFBSs reside,^2, 3^ understanding the biology of TFBS may provide insight into basic mechanisms of disease.^4^ The traditional approach to studying the function of a TFBS is through *in vitro* or *in vivo* reporter assays, where the TFBS is studied outside of its normal genomic context. Notably, few TFBSs have been modified in their native genomic milieu of the mouse and nearly all yielded imprecise mutations and genomic scarring (*e*.*g*., residual *lox*P sequence).^5-8^ Generating such mouse models with conventional embryonic stem cell targeting is labor-intensive and expensive, and the results can be uncertain given the known redundancies in TFBS utilization for target gene transcription.^9^

The emergence of clustered regularly interspaced short palindromic repeats (CRISPR) genome editing ^10, 11^ has greatly facilitated the development of precision-guided alterations to the mouse genome.^12-14^ The first generation of CRISPR editing in mice utilized three components: an endonuclease (Cas9); a single guide RNA (sgRNA) that shepherds Cas9 to the sequence to be edited; and a repair template, generally a single-strand oligodeoxynucleotide (ssODN), engineered to carry small insertions, deletions, or substitutions that are installed into the target DNA sequence during homology directed repair (HDR) of the sgRNA-Cas9 induced double-strand break.^15-17^ Three-component CRISPR successfully disrupted TFBSs in their native genomic context of mice, revealing insight into target gene expression in an in vivo setting.^18-20^ However, HDR-mediated editing is often inefficient, is limited to actively dividing cells, is associated with unwanted collateral indel mutations, and may yield off-targeting events.^21^ A second generation CRISPR platform, called base editing,^22^ was developed wherein an sgRNA directs a Cas9 nickase fused to a cytidine or adenine deaminase to target DNA for incorporation of base substitutions without the generation of a double-strand break in DNA or the need of a repair template. Base editing simplifies delivery and reduces the proportion of indels. This two-component platform edited separable TFBS in the mouse with no detectable off-targets.^23^ However, base editing is currently limited to base transitions and may generate so-called bystander substitutions at neighboring bases within the editing window, thus complicating the identification of correctly edited TFBS. Recently, a new two-component genome editing platform, called prime editing, was described in which a Cas9 nickase fused to an engineered Maloney murine leukemia virus reverse transcriptase can directly copy desired edits in the target DNA sequence from a prime editing guide RNA (pegRNA).^24^ Prime editing was demonstrated to install >175 different edits, including all possible base substitutions, in various human cell lines with limited off-target events.^24^ Thus, in principle, prime editing represents a versatile, precision-guided platform that can potentially correct all SNVs of clinical importance with limited off-targeting events.^24^ Prime editing has been reported in plants,^25-27^ in mouse embryos,^28, 29^ and in Drosophila.^30^ Here, we sought to test the efficiency of prime editing versus three-component HDR editing at a single TFBS in the mouse. We demonstrate high fidelity prime editing in germline transmitted adult mice and an unexpected phenotype in adult mice homozygous for a single base pair edit in a TFBS.

## Results

### Three-component HDR editing of a TFBS in the *Tspan2* promoter of mice

The CArG box is a TFBS for serum response factor (SRF), a widely expressed TF that directs disparate programs of gene expression.^31^ An SRF-binding CArG box is located 539 base pairs upstream of the major transcription start site of the human *TSPAN2* locus (Extended Data Fig. 1a). This CArG box is conserved in many mammalian species, including mouse (Extended Data Fig. 1b). We previously demonstrated activity of the *Tspan2* CArG box in cultured cells, but whether this TFBS is necessary for *Tspan2* expression in an animal model is unknown.^32^ To address this question, we designed an sgRNA overlapping the CArG box with the CRISPOR tool;^33^ CArG boxes are ideal TFBS for genome editing as they begin with and end in a PAM sequence (Extended Data Fig. 1b). We also designed an ssODN with three nucleotide substitutions expected to disrupt SRF binding to the CArG box (Fig. 1a). The Cas9 protein, sgRNA, and ssODN were injected into 204 mouse zygotes. Following overnight culture of injected zygotes, 143/204 (70%) viable two-cell staged embryos were transferred to 5 recipient females and 37/143 (26%) live-born births were obtained. Allele-specific PCR genotyping revealed 20/37 (54%) founder mice with evidence of correct editing, but 4/20 (20%) showed obvious indels (Extended Data Fig. 2). Due to the large number of founder mice, we selected a subset of 11/20 for on-target and off-target analyses (Extended Data Fig. 2). We validated the three base pair substitution in a founder mouse by Sanger sequencing (Fig. 1b) and bred a founder for germline transmission of the mutant CArG box. Next, heterozygous F_1_ mice were inter-crossed to generate homozygous *Tspan2* CArG mutant mice (*Tspan2*^*sg/sg*^). Normal Mendelian ratios were observed (12 *Tspan2*^*+/+*^, 27 *Tspan2*^sg/+^, and 15 *Tspan2*^sg/sg^). Several tissues were isolated from each genotype for qRT-PCR analysis. Compared to *Tspan2*^*+/+*^ controls, the expression of *Tspan2* mRNA in *Tspan2*^sg/sg^ mice was sharply attenuated in smooth muscle-rich tissues of aorta and bladder (Fig. 1c). An intermediate level of *Tspan2* mRNA expression was seen in *Tspan2*^sg/+^ mice suggesting bi-allelic expression (Fig. 1c). Although *Tspan2* mRNA in heart and brain is, respectively, similar to or more abundant than *Tspan2* in aorta (Extended Data Fig. 3), no change in expression was detected in heart or brain of heterozygous or homozygous CArG mutant mice (Fig. 1c). In situ hybridization studies confirmed loss of *Tspan2* in smooth muscle cells in blood vessels of the heart, but not surrounding cardiac tissue (Extended Data Fig. 4). Collectively, these findings demonstrate the critical role of a single TFBS for cell-specific expression of *Tspan2* in adult mice.

**Fig. 1.**
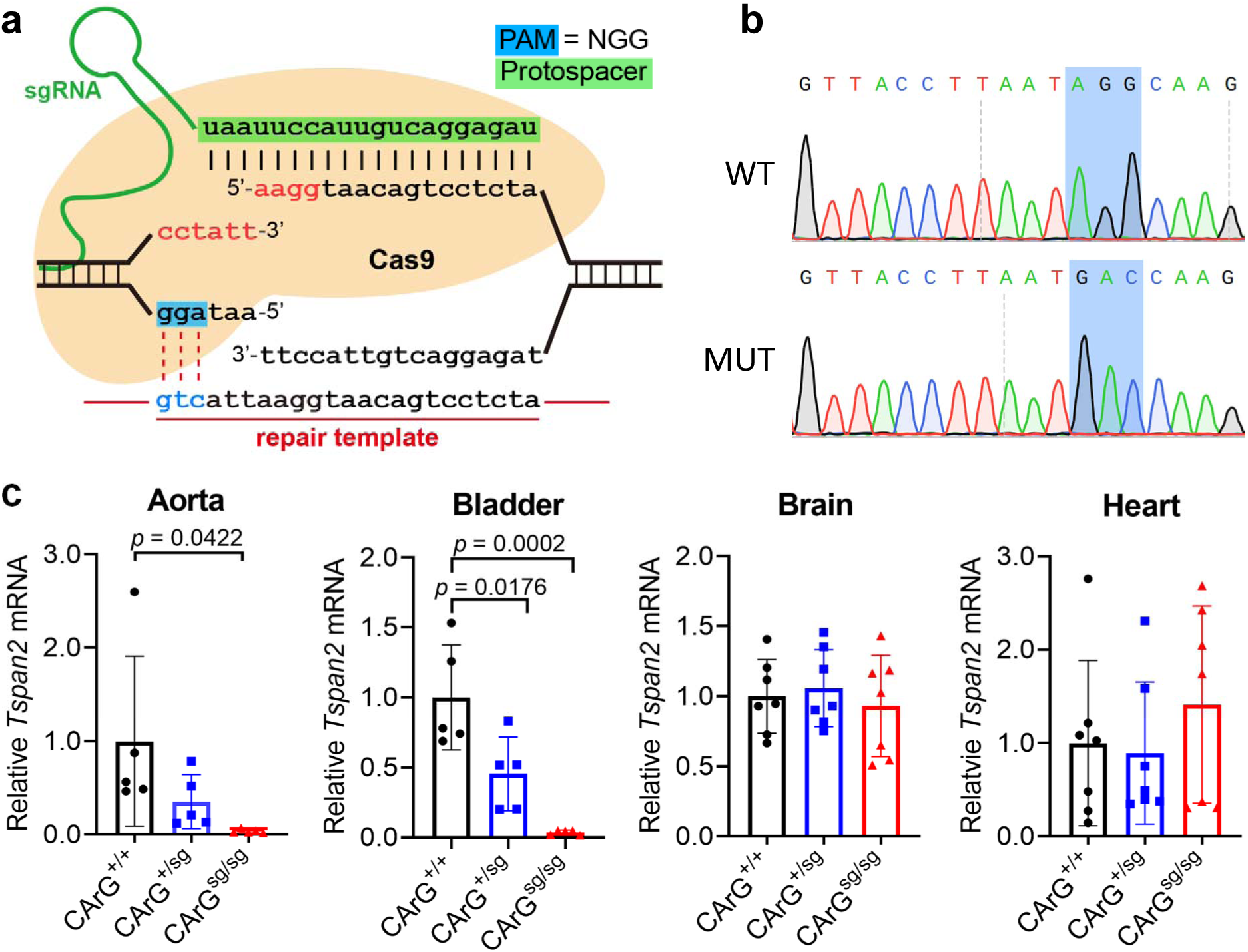
HDR-mediated editing of *Tspan2* CArG box. **a**, Targeting strategy with CArG box (red) and 3 bp substitution in blue. **b**, Sanger sequencing of CArG box showing correct edit in a mutant founder. **c**, qRT-PCR of *Tspan2* in indicated tissues and genotypes (n = 5 to 7 mice/genotype). Black, blue, and red bars here and below represent relative (wild type set to value of 1) mean *Tspan2* mRNA (± STD) in wild type, heterozygous, and homozygous genotypes. sg, sgRNA edited mouse.

### Prime editing of the *Tspan2* CArG box in mice

Inspired by previous in vitro studies demonstrating an attenuation in SRF binding to CArG boxes carrying single base pair substitutions,^34^ we set out to use the recently described prime editing platform^24^ to target the same CArG box of the *Tspan2* promoter with a single base substitution. Several versions of prime editing (PE) plasmids carrying the Cas9 nickase fused to reverse transcriptase exist, but we selected the PE2 plasmid since this version of prime editor showed low-level indels in cultured cells.^24^ Optimal in vitro transcription of pCMV-PE2^24^ was achieved by extending the incubation time to 3 hours and treating samples with RNase inhibitors (Methods and Extended Data Fig. 5). A synthetic pegRNA was generated with the following sequence features: the same 20 nucleotide protospacer sequence used for the HDR experiment followed by the scaffold extended with 10 nucleotides that correspond to the reverse transcriptase template and a 16 nucleotide primer binding site (Fig. 2a). The single base edit, a C>G transversion, was engineered at position +8 of the reverse transcriptase template (Fig. 2a). In vitro transcribed PE2 mRNA and synthetic pegRNA were injected into 234 mouse zygotes. Following overnight culture of injected zygotes, 175/223 (78%) viable two-cell staged embryos were transferred to 5 recipient females and 47/175 (27%) live-born births were obtained. Restriction digestion of the PCR product from each live-born pup revealed 12/47 (26%) founder mice with evidence of correct editing (Extended Data Fig. 6). Sanger sequencing of a founder mouse showed precise incorporation of the C>G transversion (Fig. 2b). Two of the sequence-confirmed PE2 founders transmitted the edited allele through the germline for heterozygous intercrossing. Normal Mendelian ratios were seen in F_1_ pegRNA mice (9 *Tspan2*^*+/+*^, 13 *Tspan2*^peg/+^, and 5 *Tspan2*^peg/peg^). Remarkably, the expression of *Tspan2* mRNA was nearly abolished in aorta and bladder of *Tspan2*^peg/peg^ adult mice with little change in brain and heart (Fig. 2c). Mice homozygous for either the three base pair or single base pair CArG box edit were outwardly normal and could breed.

**Fig. 2.**
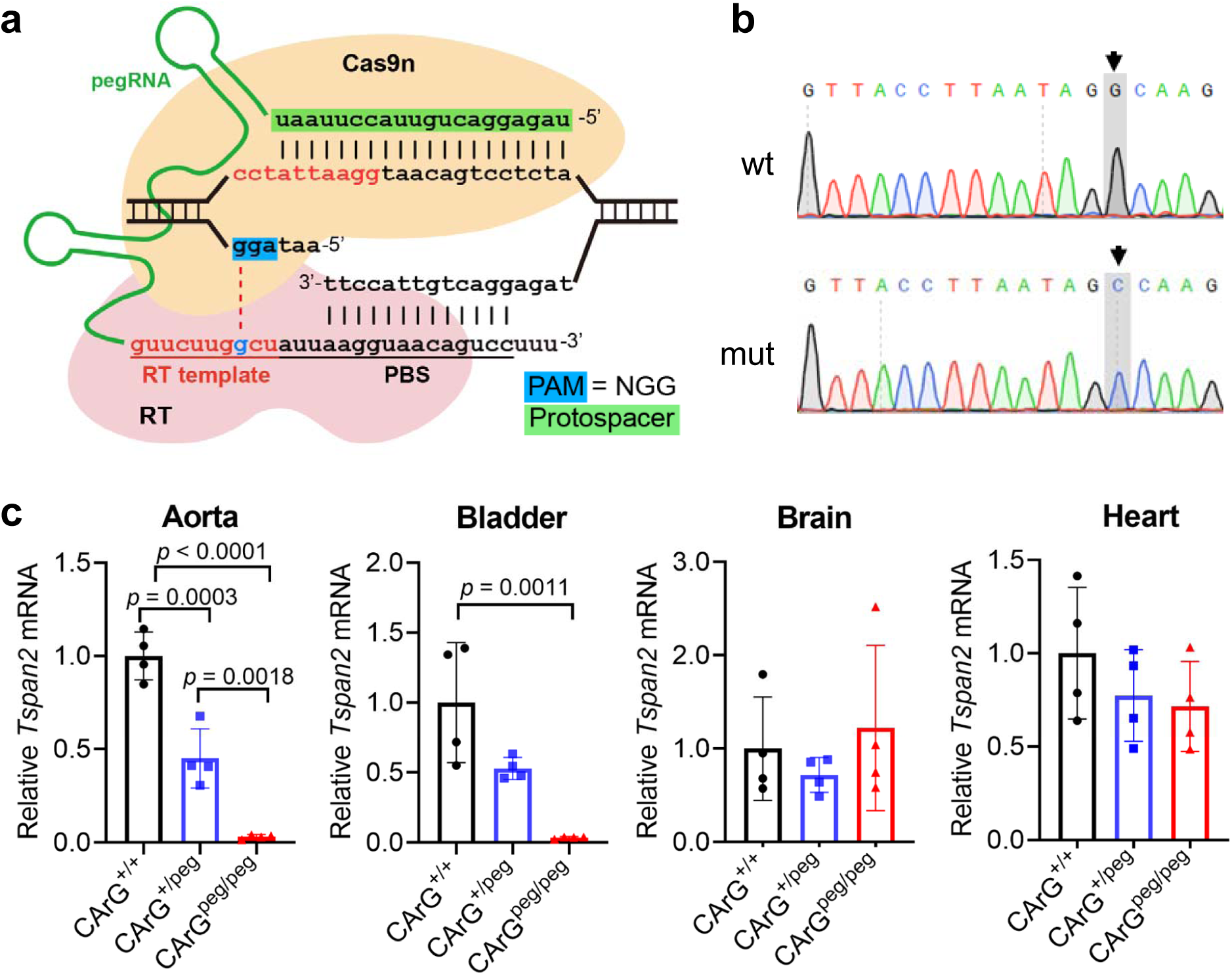
PE2-mediated editing of *Tspan2* CArG box. **a**, Targeting strategy with CArG box in red and 1 bp substitution in blue. **b**, Sanger sequencing of CArG box showing correct edit in a mutant founder. **c**, qRT-PCR of *Tspan2* in indicated tissues and genotypes (n = 4 mice per genotype). peg, pegRNA edited mouse; RT, reverse transcriptase; PBS, primer binding site.

Interestingly, a long non-coding RNA (lncRNA) overlaps the *Tspan2* locus in the mouse genome. The presumptive promoter of this lncRNA, called *Tspan2os*, is located in the first intron of *Tspan2* about 900 base pairs 3’ from the CArG box (Fig. 3a). We surmised that expression of *Tspan2os* could be similarly dependent on the targeted CArG box. Indeed, qRT-PCR revealed a notable reduction in *Tspan2os* in aorta and bladder of mice homozygous for the C>G transversion (Fig. 3b). Taken together, these findings demonstrate an essential role of a single base within a TFBS in co-regulating an mRNA-lncRNA gene pair.

**Fig. 3.**
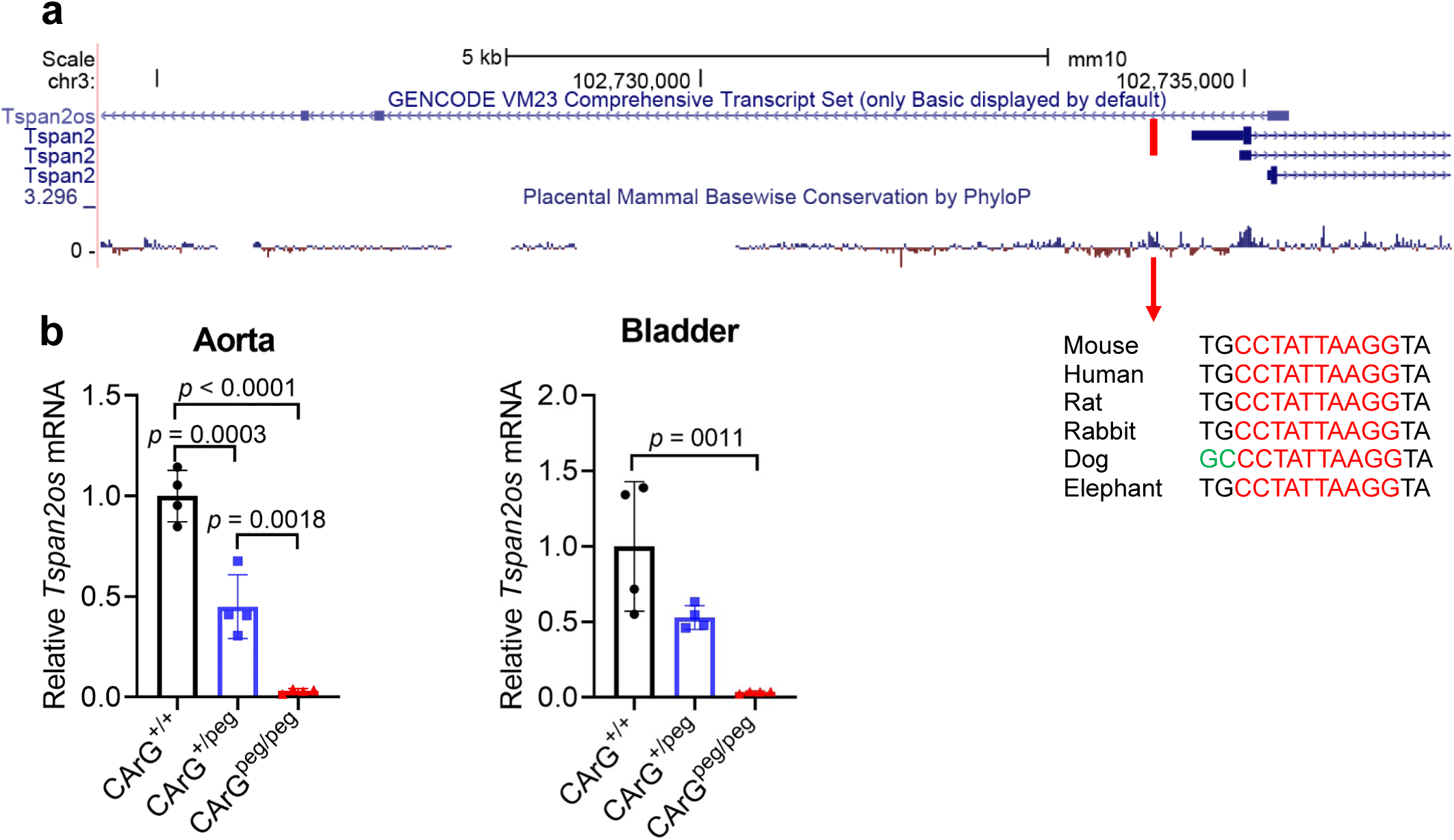
Mouse *Tspan2* and *Tspan2os* loci. **a**, UCSC Genome Browser screenshot of the 5’ mouse *Tspan2* locus and overlapping, divergently transcribed lncRNA, *Tspan2os*. The sequence of the CArG box (red line at top) shown here is the complement of that shown in Extended Data Fig. 1b due to direction of transcription in mouse versus human *Tspan2*. Note the CArG box falls within a high degree of mammalian conservation (red arrow). **b**, qRT-PCR of *Tspan2os* RNA in aorta and bladder of indicated PE2-mediated genotypes. n=4 aortae from each genotype.

### On-target editing fidelity in HDR versus prime edited founder mice

Genome editing with wild type Cas9 can elicit undesired editing outcomes such as indels among a large fraction of edited cells.^35^ On the other hand, prime editing, particularly with the single-nick PE2 system, yields a much higher purity of edited products.^24^ Accordingly, we performed targeted sequencing analysis on genomic DNA derived from the spleen of 11 HDR and 12 PE2 founder mice to evaluate the fidelity of on-target editing. The mean percentage of sequencing reads with correct on-target editing was 55.65% (range 1.67% - 95.56%) for HDR founder mice versus 20.74% (range 2.66% - 50.94%) for PE2 founder mice (t-test, *p* = 0.0067). Despite a significantly higher frequency of on-target editing, many of the HDR founders showed undesired indels (mean of 40.11%, range 0.91% - 93.91%). In contrast, none of the PE2 founders displayed indels above background at the on-target editing site (Fig. 4a, b). CRISPResso analysis further documented the frequency of indels in each of the founder mice (Fig. 4c, d). These results demonstrate precise PE2-mediated on-target editing, with no spurious indels, in the C>G transversion of the *Tspan2*/*Tspan2os* CArG box.

**Fig. 4.**
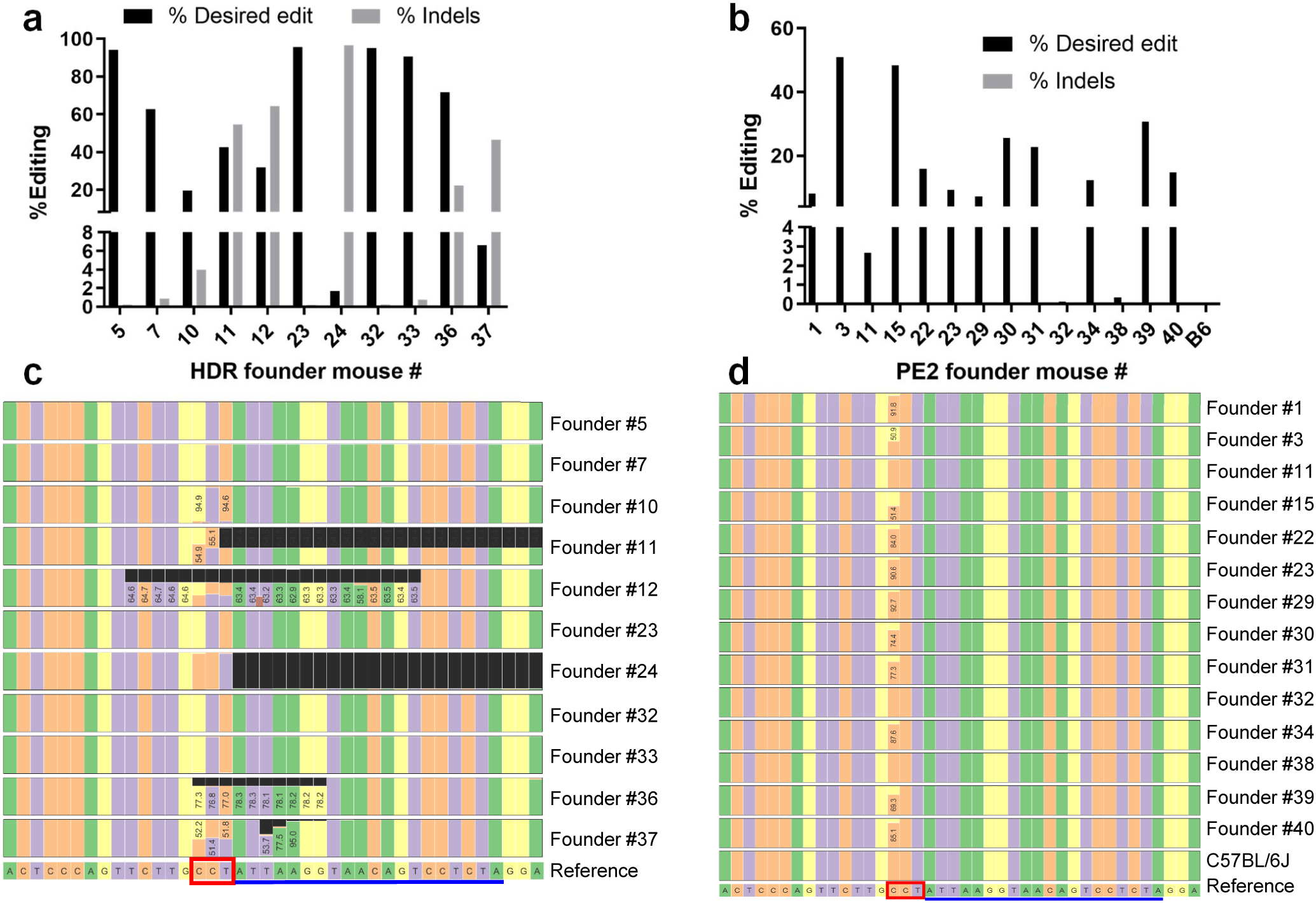
On-target sequence fidelity at the *Tspan2* CArG box. Percent editing across HDR (**a**) and PE2 (**b**) founder mice. CRISPResso sequence output for individual founders from sgRNA (**c**) and pegRNA (**d**) study. Protospacer (blue line) and PAM (red box) are indicated as are numbers indicating frequency of correct edits. Black boxes in panel **c** indicate deletions; some indels are not displayed in HDR founders due to failure of CRISPResso to align them.

### Off-target editing in HDR versus prime edited founder mice

There are >1,200 permutations of the CArG box across mammalian genomes.^36^ Because the *Tspan2*/*Tspan2os* CArG box encompasses the PAM and PAM proximal protospacer sequence, we considered the possibility of inadvertent targeting of other CArG boxes by either HDR- or PE2-mediated editing and, if present near a target gene, reduced gene expression as shown here for the *Tspan2/Tspan2os* gene pair. CRISPOR analysis of the protospacer sequence targeting the *Tspan2*/*Tspan2os* CArG box revealed specificity scores of 80 (MIT) and 93 (CFD) and 0, 1, 0, 12, and 68 predicted off-targets with 0, 1, 2, 3, or 4 mismatches, respectively. Of the 53/81 (65%) CRISPOR predicted off-targets harboring a potential SRF-binding CArG box, only 7/53 (13%) are located within four kilobases of the transcription start site where all known functional CArG boxes reside (Supplementary Table 1).^36^ To address whether off-targeting events that did not segregate upon breeding could lead to local gene repression similar to *Tspan2/Tspan2os*, we performed bulk RNA-seq analysis of aortae from *Tspan2*^*+/+*^ versus either *Tspan2*^*sg/sg*^ or *Tspan2*^*peg/peg*^ mice. This analysis revealed no significant decrease in expression of transcription units adjacent to the 81 CRISPOR predicted off-targets (Supplementary Table 1). Moreover, the only target genes significantly reduced in HDR (*Tspan2*^*sg*/sg^) and PE2 (*Tspan2*^*peg/peg*^) mice were *Tspan2* and *Tspan2os*, both of which showed ∼90% decrease versus *Tspan2*^*+/+*^ aorta (Fig. 3b, Fig. 5). Several significantly changed genes in *Tspan2*^*sg*/sg^ or *Tspan2*^*peg/peg*^ mice harbor a proximal CArG box that were not identified by CRISPOR; however, only one of these (*Hist2h2be*) was downregulated in mutant aorta (Supplementary Table 2). Of note, the two conserved CArG boxes and flanking sequence of *Hist2h2be* have 13 and 17 mismatches with the protospacer, making them unlikely targets for the sgRNA or pegRNA.

**Fig. 5.**
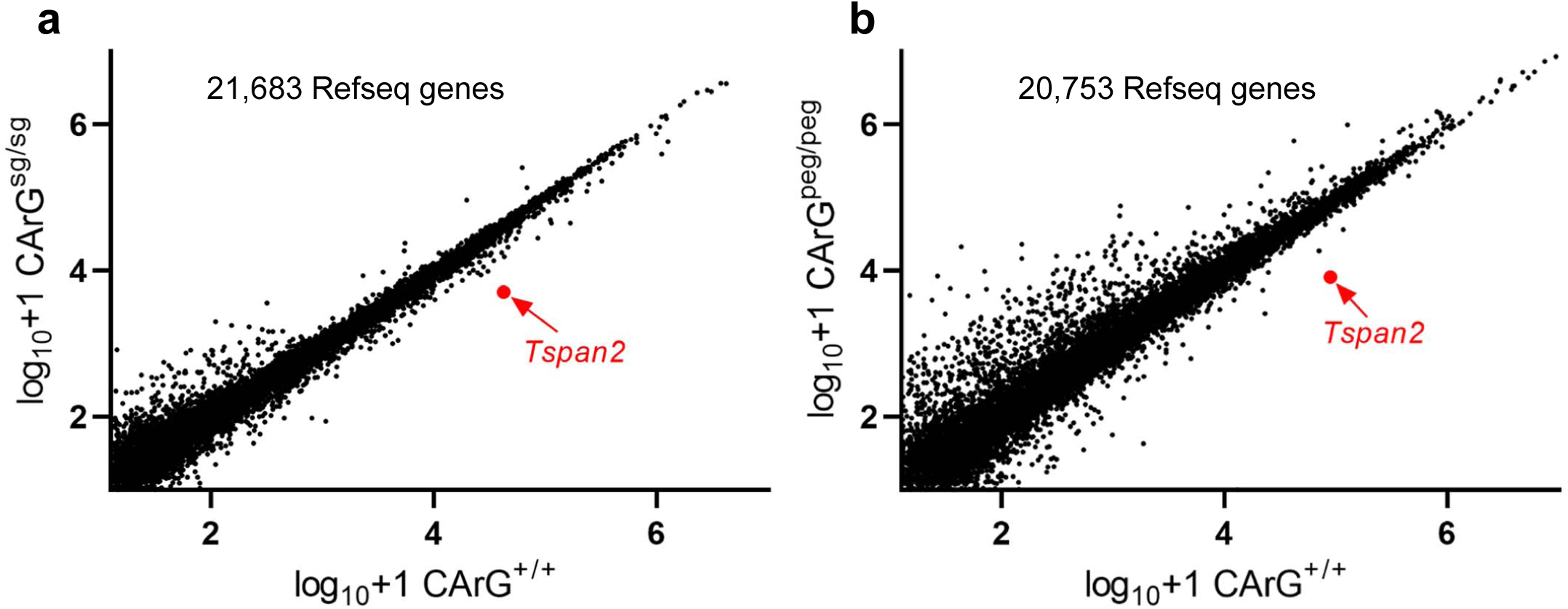
Bulk RNA-seq of aortae from HDR and PE2-edited mice. Scatter plots between HDR **a**, (sgRNA) and **b**, PE2 (pegRNA) mice. The position of differential *Tspan2* normalized reads is indicated in red. There was no overlap in genes up or down-regulated between the HDR and PE2 scatter plots. Many of the up-regulated transcripts, particularly in the pegRNA experiment, are due to large deviations in reads among single replicates. For a listing of the significantly regulated genes, please see Supplementary Table 2. n=4 aortae for each genotype.

To assess off-targeting events in HDR versus PE2 edited mice with a sensitive, unbiased genome-wide method, we performed the recently described <u>c</u>ircularization for <u>h</u>igh-throughput <u>a</u>nalysis of <u>n</u>uclease <u>g</u>enome-wide <u>e</u>ffects by <u>seq</u>uencing (CHANGE-seq) using wild type Cas9 nuclease.^37^ CHANGE-seq revealed 105 and 188 predicted off-targets for Cas9 complexed with pegRNA or sgRNA, respectively (Fig. 6a, b). 21/81 (26%) CRISPOR predicted off-targets overlapped with those derived from CHANGE-seq, and of the CHANGE-seq off-targets overlapping in both sgRNA and pegRNA samples (Fig. 6c), only 2/49 (4%) were found in the CRISPOR pool. Next, we interrogated each of the HDR and pegRNA founder mice for evidence of unintended off-target mutations at a total of 244 target sites predicted with CHANGE-seq. We performed targeted multiplex sequencing in 244 CHANGE-seq on- and off-target sites (identified in the sgRNA or pegRNA groups) and in 13 CasOFFinder^38^ sites, using rhAmpSeq, an approach we previously validated for concordance with standard targeted sequencing.^37^ We observed off-target mutations at relatively high frequencies of approximately 5.5% to 60.2% across five sites in 5 of 11 HDR founder mice (Fig. 6d). In contrast, we did not detect off-target mutations with frequencies above background in pegRNA founders or unedited control mice from the same colony (Fig. 6e). In two sgRNA founder mice, we detected off-target mutations at three distinct sites in the same animal. Off-target mutations at two off-target sites were detected in multiple sgRNA founder mice. A full listing of the frequency of off-targeting events across all CHANGE-seq targets for both HDR and PE2 founder mice can be found in Supplementary Table 3 (pool 1 targets) and Supplementary Table 4 (pool 2 targets)..

**Fig. 6.**
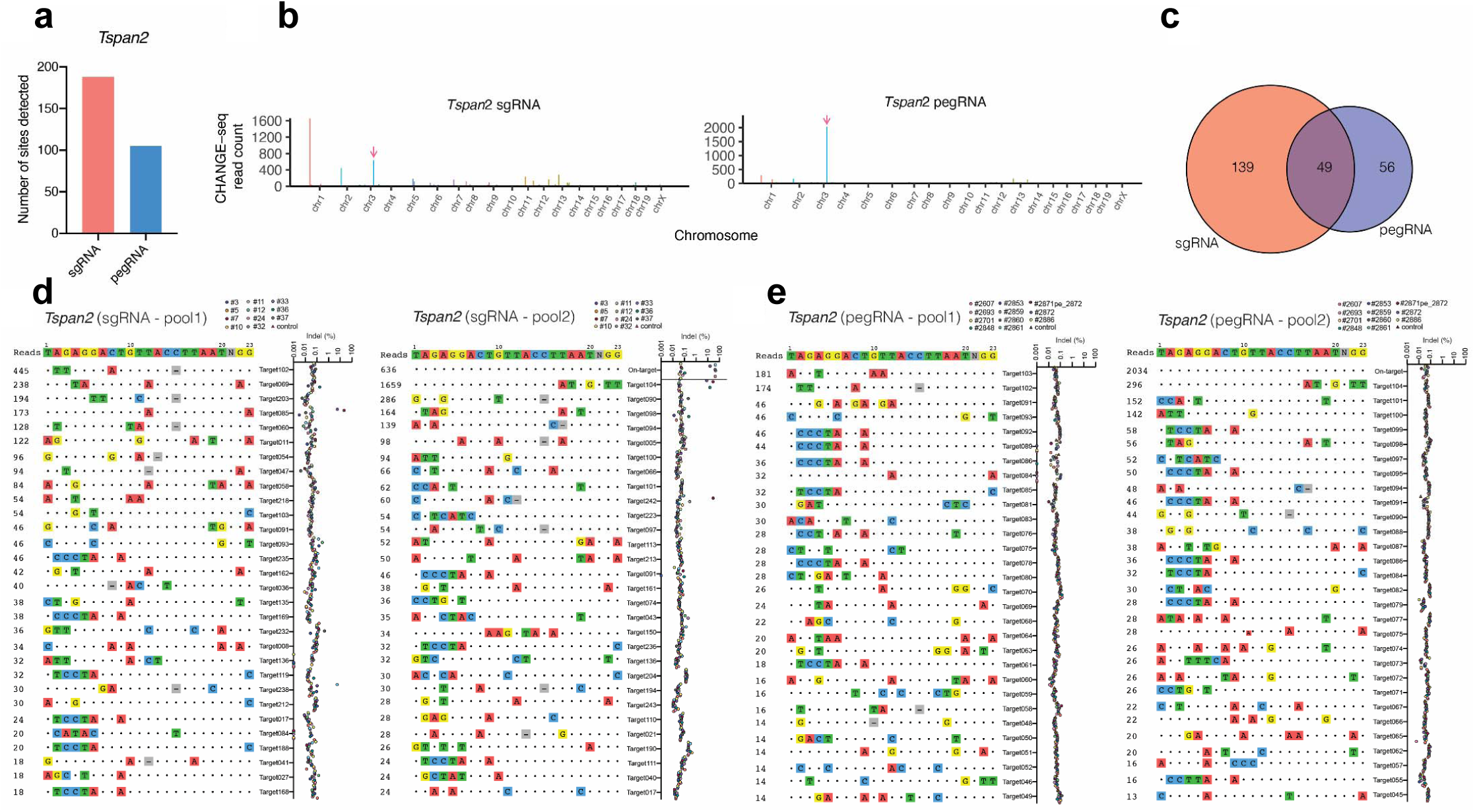
Genome-wide off-target analysis of HDR and PE2 edited founder mice. **a**, Bar plot of number of CHANGE-seq sites detected using Cas9 WT and synthetic sgRNA or pegRNA targeting *Tspan*2, on WT genomic DNA from same strain of mice used in HDR and PE2 editing experiments. **b**, Manhattan plots of CHANGE-seq detected on- and off-target sites organized by chromosomal position, for sgRNA and pegRNA, with bar heights representing CHANGE-seq read count. Arrow indicates the on-target site. **c**, Venn diagram depicting common predicted off-target sites for sgRNA (orange) and pegRNA (blue) groups. **d, e**, Indel frequencies evaluated by rhAmpSeq at on- and off-target sites detected by CHANGE-seq for sgRNA founders **d**, and for pegRNA founders **e**.

We next analyzed whether any of the 244 CHANGE-seq sites contain a CArG box. There are 66/105 (63%) and 109/188 (58%) CArG boxes in the pegRNA and sgRNA CHANGE-seq identified loci, respectively. The vast majority of these sites (171/175 or 97.7%) are distal (>4kb) from any annotated transcript. Nevertheless, we assessed whether the nearest transcription unit in these distal sites exhibited changes in bulk RNA-seq of aorta; none showed a significant decrease in expression (Supplementary Tables 5 and 6). Only 4/175 (2.3%) CHANGE-seq targets have a CArG box in close proximity to the transcription start site, but none showed any change in bulk RNA-seq of aorta (Supplementary Tables 5 and 6). Taken together, these analyses demonstrate that while there were more off-targeting events in the HDR founders (5/11 or 45%) none resulted in a CArG box-dependent decrease in gene expression.

## Discussion

The initial reporting of PE2 demonstrated versatility of editing in cultured cells with minimal off-targeting events.^24^ It is essential, however, to extend findings to more complex model systems and compare the relative efficiency of PE with other genome editing platforms such as base editing and HDR-mediated editing. Previous work in mouse embryos has shown variable fidelity of prime editing, depending on the prime editing system used.^28, 29^ However, no studies have yet to compare prime editing with conventional HDR in adult mice and there have been no reported mouse phenotypes linked to prime editing. Here, we compared the prime editing 2 (PE2) system^24^ with conventional HDR-mediated editing in adult mice using the same protospacer targeting a TFBS (CArG box) located in the *Tspan2* promoter region. *Tspan2* encodes for a membrane-associated protein highly enriched in vascular and visceral smooth muscle cell-containing tissues.^32^ We show that while both editing platforms successfully installed the engineered nucleotide substitutions within the CArG box, PE2 did so without measurable on-target indels and with low off-targeting events. HDR-mediated editing with a three-base substitution in the CArG box resulted in a similar reduction of *Tspan2* expression; however, the majority of founder mice exhibited on-target indels and several exhibited off-target editing events. Remarkably, we found that prime editing of a single nucleotide within the CArG box resulted in ∼90% reduction in expression of *Tspan2* in vascular and visceral smooth muscle tissues of adult mice; no change in expression was observed in heart muscle or in brain. A similar reduction in RNA expression was observed for an overlapping long noncoding RNA (*Tspan2os*). To our knowledge, this is the first formal demonstration of the essentiality of a single base within a TFBS for tissue-restricted gene expression in mice and the co-regulation of an mRNA/lncRNA gene pair. The severe attenuation of *Tspan2* expression in aorta and bladder provides a unique opportunity to elucidate its function in smooth muscle without the need for engineering complex Cre/*lox*P mice. Moreover, since the coding sequence of *Tspan2* was unadulterated, future genetic rescue studies could be simplified using CRISPR activation^39^ to override the regulatory edit in the CArG box. One caveat of this approach, however, relates to mRNA/lncRNA gene pairs where disruption of a shared TFBS could confound interpretation of phenotypes. Accordingly, characterization of the mice reported here will require genetic complementation studies where either the TSPAN2 protein or the *Tspan2os* long noncoding RNA are reconstituted to better interpret phenotypes. Based on natural genetic variation and in vivo mutagenesis screening in different mouse strains,^40^ we surmise there will likely be more examples of single base edits in a TFBS resulting in a significant decrease in target gene expression in mice.

The reduction of *Tspan2* in smooth muscle tissues with a single base edit to the CArG box was unexpected given the heterogeneity of SRF-binding CArG boxes across the genome.^36^ This suggests that some substitutions across the CArG box are intolerant for SRF binding, a notion supported by atomic structure studies.^41^ Interestingly, despite the abundant expression of SRF in brain^42^ and heart,^43^ the CArG box mutants generated here had no effect on expression of *Tspan2* in these tissues. This would imply that separable TFBS recognized by distinct transcription factors drive expression of *Tspan2* in heart and brain. Future studies could determine whether subtle editing of targeted TFBS around the *Tspan2* locus confer loss of expression in brain or heart.

Nearly all of our genome is noncoding, comprising tens of thousands of noncoding RNAs and millions of TFBS.^3, 44^ Most SNVs associated with disease reside in noncoding sequence but causality of such regulatory SNVs in disease is notably lacking, especially in the complex milieu of an animal model.^3^ For example, 146/164 (89%) coronary artery disease risk alleles harbor noncoding SNVs but, with the exception of the noncoding SNV near *SORT1*,^45^ we lack insight into the functional consequence of such sequence variants *in vivo*. While no known SNV exists in the *Tspan2/Tspan2os* CArG box, the large number of such TFBS^34^ would suggest the presence of potentially important CArG-SNVs that could be easily modeled with PE2 editing in the mouse. An important goal therefore will be to map all CArG-SNVs in the human genome and filter those of possible clinical relevance for further study.

A limitation of genome editing in animal model systems and in future clinical trials to correct disease-causing mutations is unintended editing at the desired editing site or at distal off-targets. Whole genome sequencing and screening experiments in animal models have demonstrated low-level off-targeting events with wild type Cas9^46-49^ and, more recently, prime editing^28^ though it must be stressed that these studies were necessarily limited to very few guide RNAs. A broader analysis in mouse and rat showed measurable off-targeting in 23% of Cas9-sgRNA experiments, with most off-targeting nearly eradicated by high-fidelity Cas9 nucleases.^50^ Here, we found essentially no on-target indels above the limits of detection in 12 PE2 founder mice whereas unwanted on-target indels were detected in the majority of HDR founders. We also conducted a genome-wide analysis using the recently developed CHANGE-seq protocol^37^ and identified nearly two-fold greater number of candidate off-target sites with wild type Cas9 and sgRNA than wild type Cas9 and the pegRNA. The reason for the variance in candidate off-targets is unclear but may be due to technical reasons related to the different structures of the sgRNA and pegRNA. We interrogated 244 candidate off-target sites in all founder mice and found off-targeting events in 5/11 (45%) HDR founders. In contrast, we did not detect evidence of off-target mutations in PE2-edited founder mice. Although the majority of off-targets predicted by CHANGE-seq (and CRISPOR) harbor a CArG box, bulk RNA-seq of aortic tissue failed to reveal reduced expression of any associated transcript though it is possible transcripts could be reduced in other tissues not evaluated here. Moreover, since bulk RNA-seq occurred in the F_2_ generation, it is possible any distal edit of a CArG box resulting in gene repression would have been lost following segregation of alleles during meiosis.

In summary, we provide the first comparison of prime editing with HDR-mediated editing in adult mice and show the PE2 system is effective in the installation of a single nucleotide substitution within a TFBS without on-target indels or detected off-targeting events. This single base replacement confers a near complete loss in expression of the *Tspan2*/*Tspan2os* gene pair in smooth muscle-rich tissues, allowing for future characterization of phenotypes under baseline and stress-induced conditions. Under the experimental conditions reported here, the PE2 platform yielded precision-guided editing without unwanted mutations. These desirable attributes as well as the development of computational tools for optimal pegRNA design,^51^ should stimulate additional comparative studies to further assess the fidelity of prime editing in mice. Finally, it will be of great interest to assess the potential utility of the prime editing platform in somatic editing of both prenatal and postnatal animal models.

## Methods

### HDR-mediated genome editing of *Tspan2* CArG box in mice

The mouse experiments in this study were approved by local institutional animal care and use committees at Cornell University (#2000-0122) and Medical College of Georgia at Augusta University (#2019-0999 and #2019-1000). Fertilized oocytes derived from male B6(Cg)-Tyr^2J^ /J (Jackson labs, stock #000058) and superovulated female FVB/NJ (Jackson labs, stock #001800) mice were microinjected with 50 ng/μl of wild type Cas9 mRNA (TriLink Biotechnologies, San Diego, CA), 50 ng/μl of sgRNA (Synthego Corp., Menlo Park, CA), and 25 ng/μl of ULTRAmer Standard Desalting single-strand ODN (Integrated DNA Technologies, Coralville, IA) harboring a CCT>GTC substitution in the *Tspan2* CArG box (see Figure 1A and Extended Data Fig. 2). All CRISPR components were dissolved in nuclease-free water (Ambion #9932) and diluted in injection buffer (100 mM NaCl; 10 mM Tris-HCl, pH, 7.5; and 0.1 mM EDTA) and injected into the pronucleus and cytoplasm using a Nikon Eclipse TE200 microscope equipped with Eppendorf FemptoJet 4x, Eppendorf TransferMan NK manipulator, and Eppendorf CellTram Air vacuum (Enfield, CT). Injected eggs were cultured in KSOM medium at 37^°^ C overnight and viable two-cell staged embryos were transferred to the oviducts of pseudopregnant female mice of strain B6D2F1/J (Jackson labs, stock #1000006) and allowed to develop to full term. Founder mice were weaned 21 days post-parturition and genomic DNA from tail snips of founder mice isolated with Gentra Puregene Tissue Kit (Qiagen Sciences #158667; Germantown, MD) according to manufacturer’s instructions. Allele-specific ODN primers (Integrated DNA Technologies, Coralville, IA) were used to PCR genotype each founder pup for the presence of *Tspan2* CArG box editing (see Extended Data Fig. 2). ODNs for HDR-mediated repair and PCR genotyping are listed in Supplementary Table 7. Selected HDR founder mice were bred to strain C57BL/6J mice (Jackson labs, #000664) to pass the CCT>GTC substituted CArG box allele through the germline for heterozygous intercrossing and gene expression analysis. In addition, 11 HDR founders were analyzed for on-target and off-target editing as described below.

### Prime editing of *Tspan2* CArG box in mice

The same strains of mice used in HDR-mediated editing were used for prime editing. The Cas9 nickase-reverse transcriptase plasmid (pCMV-PE2, Addgene #132775, Watertown, MA) was linearized with PmeI (New England Biolabs #R0560S, Ipswich, MA) for 3 hours at 37°C, excised from an agarose gel, purified with a Monarch^®^ DNA Gel Extraction kit (New England Biolabs #T1020S, Ipswich, MA), and incubated with RNAsecure™ RNase Inactivation Reagent (ThermoFisher Scientific #AM7005, West Columbia, SC) for 15 minutes at 65°C. Linearized and purified pCMV-PE2 was then in vitro transcribed using mMESSAGE mMACHINE & Ultra kit (ThermoFisher Scientific #AM1345, West Columbia, SC) for 3 hours at 37°C in the presence of RNasin Ribonuclease Inhibitor (Promega #N2111, Madison, WI), and PE2 mRNA was purified with MEGAclear™ Transcription Clean-Up kit (ThermoFisher Scientific, #AM1908, West Columbia, SC) according to the manufacturer’s instructions. The prime editing guide RNA (pegRNA) was synthesized using Synthego’s CRISPRevolution platform with solid phase phosphoramidite chemistry. Based on the original prime editing report,^24^ we selected a reverse transcriptase (RT) template of 10 nucleotides in length, inclusive of the C>G transversion, and a primer binding site (PBS) of 16 nucleotides in length (see Figure 2A). Three 2’-O-methyluridinylates were attached at the 3’ end of the PBS (see Figure 2A) and stabilized with phosphorothioate backbones. The first three bases of the protospacer were modified as 2’-O-methyl derivatives and stabilized as phosphorothioates. The pegRNA was purified using reversed-phase high-performance liquid chromatography (Buffer A, 0.1M TEAA; Buffer B, 50% 0.1M TEAA/50% acetonitrile, 15%-95% B gradient in 15 minutes), and their identities were confirmed using an Agilent 1290 Infinity II liquid chromatography system coupled with Agilent 6530B Quadrupole time-of-flight mass spectrometry (Agilent Technologies, Santa Clara, CA) in a negative ion polarity mode. Pronuclear/cytoplasmic injections were carried out with 25 ng/μl each of the PE2 mRNA dissolved in RNAse-free water and the synthetic pegRNA dissolved in nuclease-free water and diluted in the same injection buffer used for HDR editing. Genomic DNA from tail snips of founder mice was isolated with Gentra Puregene Tissue Kit (Qiagen Sciences #158667; Germantown, MD) according to manufacturer’s instructions and PCR genotyped with primers flanking the CArG box followed by restriction digestion of the PCR amplicon with PflMI (Van9lI). The latter restriction site (CCA[N]_5_TGG) would be generated with installment of the C>G transversion (see Extended Data Fig. 5). ODNs for PE2-mediated repair and PCR genotyping are listed in Supplementary Table 7. Selected PE2 founder mice were bred to strain C57BL/6J mice (Jackson labs, #000664) to pass the C>G transversion allele through the germline for heterozygous intercrossing and gene expression analysis. In addition, 12 PE2 founders were analyzed for on-target and off-target editing as described below.

### Sanger sequencing

Initial PCR genotyping informed us of founder mice carrying either the three base-pair substitution (HDR) or the single base substitution (PE2) in the *Tspan2* CArG box (see Extended Fig. 2 and 5, respectively). PCR amplicons from several of these founders were prepared for cloning into the pCR4-TOPO TA vector (ThermoFisher Scientific #450071) and plated on LB agar plates for ampicillin resistant colony isolation, purification, and PCR validation with original primers to ensure the presence of a clone. Several independent clones from each founder analyzed were then prepared for Sanger sequencing (GENEWIZ^®^, Research Triangle Park, NC). Representative electropherograms were cropped in Adobe Photoshop for presentation.

### Quantitative RT-PCR analysis

Indicated tissues from adult mice were rapidly excised, cleaned of adhering tissue in ice cold phosphate buffered saline, and plunged in liquid nitrogen. Tissues were homogenized with a Minilys homogenizer (Bertin Technologies, Rockville, MD) using a Precellys Lysing Kit (VWR Scientific, Radnor, PA). Total RNA was extracted from thoroughly homogenized tissues via miRNeasy Mini Kit (#217004, Qiagen) according to the manufacturer’s directions. The concentration of RNA was measured by a Nanodrop 2000 spectrophotometer (ThermoFisher Scientific), and 200-500 nanograms of total RNA was programmed for cDNA synthesis using an iScript™ cDNA Synthesis Kit (#1708890 Bio-Rad; Hercules, CA). Universal SYBR Green Supermix (Bio-Rad)-based qRT-PCR was carried out in a CFX386 Touch™Real-Time PCR Detection System (Bio-Rad). *Tspan2* mRNA and *Tspan2os* RNA expression were calculated by the 2^-ΔΔCt^ method using *Hprt* as an internal housekeeping control. Primer sequences used in this study are listed in Supplementary Table 7. Expression data were derived from tissues of independent mice (sample sizes indicated in figure legends) of each genotype and each data point in the scatter plots represents the mean of technical replicates (n=3) for each mouse.

### Immuno-RNA fluorescence in situ hybridization assay

*Tspan2*^*+/+*^ and *Tspan2*^*sg/sg*^ heart and aorta were fixed in 10% neutral buffered formalin, paraffin embedded, and cut at 5 microns. Sections were processed for combined immunofluorescence of LMOD1 protein (Proteintech, #15117-1-AP; 1:200 dilution) and RNA in situ hybridization of *Tspan2* mRNA (RNAscope, ACD) according to the manufacturer’s instructions. Alexa fluor 488 secondary antibody was used to detect LMOD1 protein (Thermofisher). Signals were obtained with a LSM 900 confocal laser scanning microscope (Zeiss) using the Zeiss Blue software system for image acquisition and processing.

### Targeted sequencing analysis for on-target editing efficiency

Genomic DNA was isolated from the spleen of indicated HDR and PE2 founder mice by DNeasy Blood & Tissue Kit (QIAgen, #69504; Germantown, MD). Primers, with adapters for barcoding, were used to amplify 288 base pairs around the CArG box (Supplementary Table 7). 0.5 μL of PCR product was used as a template in a barcoding PCR reaction, consisting of two minutes at 98°C, followed by 10 cycles of denaturation for 10 seconds at 98°C, annealing for 20 seconds at 61°C, extension for 30 seconds at 72°C, and a final 2 minute extension at 72°C. Barcoded products were pooled and gel purified from a 1% agarose gel using a QIAgen kit (#28115) to remove primers before quantitation with a Qubit dsDNA HS Kit (ThermoFisher Scientific #Q32851). Samples were loaded onto an Illumina MiSeq instrument with a 300 cycle v2 kit for sequencing. Greater than 30,000 reads were collected for each sample. Sequencing reads were demultiplexed using the MiSeq Reporter (Illumina) and fastq files were analyzed using Crispresso2 Batch Analysis.^52^ An analysis window of 10 was used to identify indels. Analysis of some nuclease-treated mice yielded substantial fractions of non-aligning reads (1-79% of total reads). Visual inspection of these sequences in Geneious DNA analysis software (Biomatters Inc., San Diego, CA) indicated that they harbored larger (>10nt) deletions, so non-aligning reads were added to the quantified indels. Non-aligning reads were less than 1% of total reads for all prime-editor treated mice.

### CHANGE-seq analysis for off-target events

Genomic DNA from B6(Cg)-Tyr^2J^ /J (Jackson labs, stock #000058) and FVB/NJ (Jackson labs, stock #001800) mouse liver was purified using Gentra Puregene Kit (Qiagen) according to manufacturer’s instructions and combined for CHANGE-seq as previously described.^37^ Briefly, genomic DNA was tagmented with a custom Tn5-transposome to an average length of 400 bp, followed by gap repair with Kapa HiFi HotStart Uracil+ DNA Polymerase (KAPA Biosystems) and Taq DNA ligase (#M0208, New England Biolabs). Gap-repaired tagmented DNA was treated with USER enzyme (#M5508, New England Biolabs) and T4 polynucleotide kinase (#M0201, New England Biolabs). Intramolecular circularization of the DNA was performed with T4 DNA ligase and residual linear DNA was degraded by a cocktail of exonucleases containing Plasmid-Safe ATP-dependent DNase (Lucigen #E3101K), Lambda exonuclease (#M0262, New England Biolabs) and Exonuclease I (#M0293, New England Biolabs). *In vitro* cleavage reactions were performed with 125 ng of exonuclease-treated circularized DNA, 90 nM of SpCas9 protein (#M0386, New England Biolabs), NEB buffer 3.1, and 270 nM of sgRNA or pegRNA, in a 50 μL volume (please note: there is no purified Prime Editor protein at this time). Cleaved products were A-tailed, ligated with a hairpin adaptor (New England Biolabs), treated with USER enzyme (New England Biolabs) and amplified by PCR with barcoded universal primers NEBNext Multiplex Oligos for Illumina (#E7335, New England Biolabs), using Kapa HiFi Polymerase (KAPA Biosystems). Libraries were quantified by qPCR (KAPA Biosystems) and sequenced with 151 bp paired-end reads on an Illumina MiniSeq instrument. CHANGE-seq data analyses were performed using open-source CHANGE-seq analysis software (https://github.com/tsailabSJ/changeseq).

### Targeted sequencing by rhAmpSeq and indel analysis

On- and off-target sites for sgRNA and pegRNA targets were amplified from mouse spleen genomic DNA using two pools of customized rhAMPSeq libraries (Integrated DNA Technologies, Coralville, IA) (primers available upon request). Sequencing libraries were generated according to manufacturer’s instructions and sequenced with 151 bp paired-end reads on an Illumina NextSeq instrument. Indel analyses were conducted using custom Python code and open-source bioinformatic tools. First, paired-end high-throughput sequencing reads were processed to remove adapter sequences with trimmomatic (version 0.36),^53^ merged into a single read with FLASH (version 1.2.11)^54^ and mapped to mouse genome reference mm10 using BWA-MEM (version 0.7.12).^55^ Reads that mapped to on-target or off-target sites were realigned to the intended amplicon region using a striped Smith–Waterman algorithm as implemented in the Python library scikit-bio; indels were counted and reported with total read counts.

### Bulk-RNA-seq

RNA-seq and data analysis was carried out in the Genome Research Center at the University of Rochester School of Medicine and Dentistry. Total RNA from individual aortae cleaned of periadventitial tissue was extracted and quantitated as described above and RNA quality assessed with the Agilent Bioanalyzer (Agilent, Santa Clara, CA). TruSeq-Stranded mRNA Sample Preparation Kit (Illumina, San Diego, CA) was used for next generation sequencing library construction per manufacturer’s protocols. Briefly, mRNA was purified with oligo-dT magnetic beads and fragmented for first-strand cDNA synthesis with random-hexamer priming followed by second-strand cDNA synthesis using dUTP incorporation. End repair and 3’ adenylation was performed on the double-stranded cDNA and Illumina adaptors were ligated, purified by electrophoresis, and PCR-amplified with primers to the adaptor sequences to generate amplicons of ∼200-500 base pairs. Amplified libraries were hybridized to the Illumina flow cell and single-end reads of 75 nucleotides were generated using Illumina’s NextSeq550 sequencer (San Diego, CA). Raw reads were demultiplexed using bcl2fastq version 2.19.1. Quality filtering and adapter removal were performed using FastP (version 0.20.0) and cleaned reads were then mapped to *Mus musculus* (GRCm38 + Gencode-M22 Annotation) using STAR_2.7.0f. Gene level read quantification was derived using the subread-1.6.4 package (featureCounts) with a GTF annotation file (Gencode M22). Differential expression analysis was performed using DESeq2-1.22.1 with a p-value threshold of 0.05 within R version 3.5.1 (https://www.R-project.org/). PCA plot, heatmap, and Gene Ontology analysis are available upon request. RNA-seq data have been deposited in the GEO database under accession number GSE158388. Scatter plots for log_10_ +1 transformed reads of >1 were generated in Excel for wild type control *Tspan2* CArG versus HDR-edited or PE2-edited *Tspan2* CArG box.

### Analysis of off-targets for CArG box and gene expression change by RNA-seq

Predicted off-targets from CRISPOR^33^ and CHANGE-seq were interrogated for the presence of consensus CArG boxes (CCW_6_GG) or CArG-like boxes (consensus CArG box with 1 nucleotide substitution) and evidence of SRF-binding using data from ENCODE on the UCSC Genome Browser.^56^ All predicted off-target sequences were then analyzed for the nearest transcription unit and these genes were cross-referenced to the RNA-seq data for changes in RNA expression.

### Statistics and data availability

All statistical analyses were conducted in GraphPad 8.0. We tested group values for normality with the Kolmogorov-Smirnov and Shapiro-Wilk tests. Differences in means (± standard deviation) were computed either with unpaired t-test for two comparisons or one-way ANOVA followed by Tukey’s post-hoc testing for more than two comparisons. Statistical significance was assumed with a probability value of *p* < 0.05. All data generated or analyzed during this study are included in this published article and its supplementary information files or deposited in a public database (GSE 158388).

## Acknowledgments

We thank the Genomics Research Core at the University of Rochester for bulk RNA-seq and analysis. XL and JMM thank members of the Long and Miano labs for providing feedback over the course of the studies and the generous support of the Medical College of Georgia at Augusta University. We also thank Talisha Davis and Shobha Yerigenahally for managing the mouse strains and Bart Bryant and Ajay Kumar for providing comments on the manuscript. Research reported in this publication was supported by the National Institutes of Health under award numbers R01HL138987, R01HL136224, and R01HL147476 (to JMM); R01HL122686 and R01HL139794 (to XL); U01AI142756, UG3AI15055101, UG3TR002636, RM1HG009490, and R35GM118062 (to DRL who also gratefully acknowledges support from HHMI); GAN is supported by a Helen Hay Whitney postdoctoral fellowship; WZ is supported by American Heart Association Career Development Award 18CDA34110319. The content is solely the responsibility of the authors and does not necessarily represent the official views of the National Institutes of Health.

## Author contributions

PG and MC characterized the HDR-mediated CArG mutation mouse line. PG, MC, and WZ prepared tissues for RNA isolation and analysis by qRT-PCR as shown in Fig. 1c, 2c, and 3b and Extended Data Fig. 3. OJS performed RNAscope and generated images in Extended Data Fig.4. ARG optimized conditions for IVT of the prime editor and purified the final product for microinjections as shown in Extended Data Fig. 5. ARG developed and validated the PE2 genotyping assay. PG, QL, WZ, and ARG performed genomic DNA preparation, genotyping, or Sanger sequencing as shown in Fig. 1b and 2b and in Extended Data Fig. 2 and 6. PG prepared all the RNA samples for RNA-seq. QL prepared genomic DNA and PCR for the targeted sequencing studies. GAN and DRL performed the MiSeq runs and data analysis as shown in Fig 4. CRL and SQT did the CHANGE-seq and rhAmpSeq analysis as shown in Fig. 6 as well as Supplementary Tables 3-6. RJM and CMA, under the direction of JCS, performed the microinjections to generate founder mice, isolated tissues for genomic DNA purification, bred founders for germline transmission, and arranged for shipment of mice to Augusta. QL prepared the schematics to Fig. 1a and 2a. JMM and XL generated the UCSC Genome Browser data as shown in Fig. 3a and Extended Data Fig. 1. JMM generated the scatter plots in Fig. 5 and analyzed CRISPOR, CHANGE-seq, and rhAmpSeq data for CArG boxes and changes in gene expression as shown in Supplementary Tables 1, 5, and 6. XL prepared the differential RNA expression data of Supplementary Table 2. KH, JAW, and APK oversaw the synthesis of the sgRNA and pegRNA. JMM and XL assembled the teams and conceived the general design of the study. JMM oversaw the entire project and wrote the manuscript with input from all other authors.

---

**Extended Data Fig. 1.**
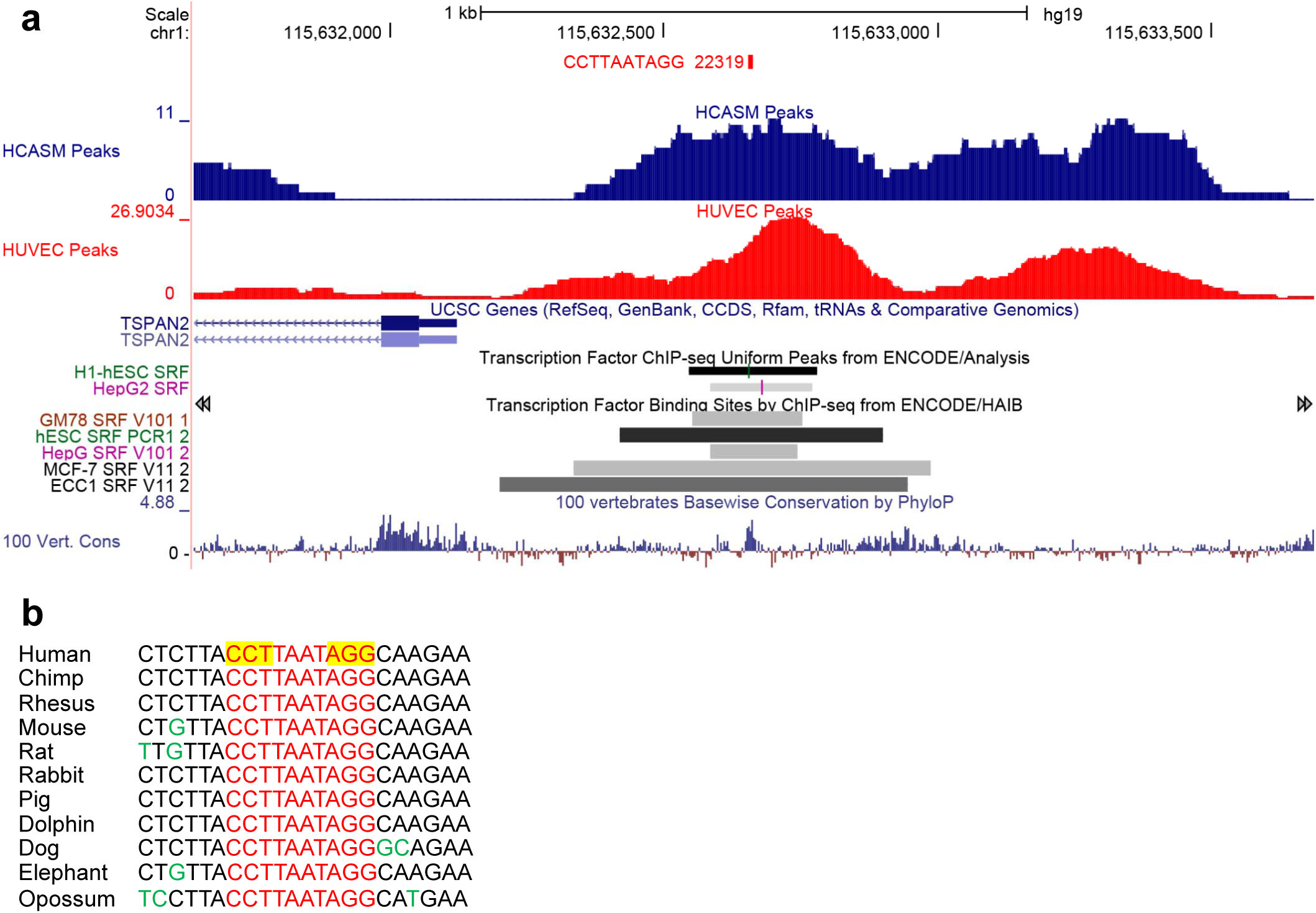
Human *TSPAN2* locus and CArG sequence. (**a**) Screenshot of UCSC Genome Browser showing CArG sequence in 5’ promoter region (red at top) and ChIP-seq data for SRF-binding in human coronary artery SMC (blue) and human umbilical vein endothelial cells (red). Also shown are SRF ChIP-seq data from ENCODE (dark bars). Note the *TSPAN2* locus is transcribed from the Crick strand in this view. There is no annotated lncRNA associated with the *TSPAN2* locus in human (see Fig. 3a for mouse lncRNA,*Tspan2os*). (**b**) Conservation of CArG sequence (red) is shown to left with PAM sequences highlighted yellow; he PAM sequence utilized in this study is to left. Sequence divergence flanking CArG box is indicated with green nucleotides.

**Extended Data Fig. 2.**
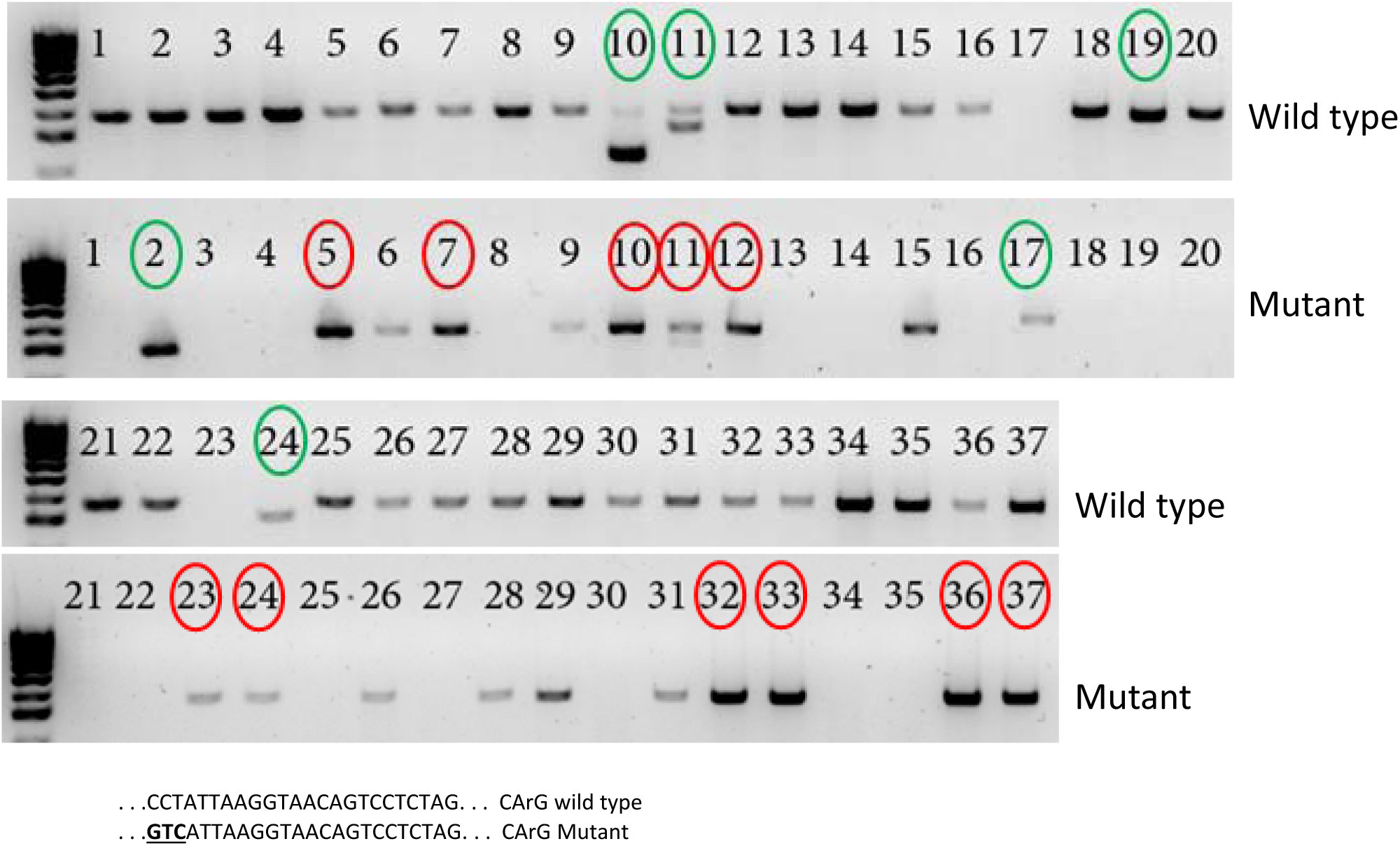
Genotyping of original founder mice derived from HDR editing of *Tspan2* CArG box. Primers specific to the wild type sequence or the 3 base pair substitution (CCT > GTC) were used in separate PCR reactions to generate the indicated bands above. Those founders circled in green indicate the presence of obvious indels. Those founders circled in red represent the mutants analyzed for further study. Sequences below represent wild type and mutant (bold underlined) CArG boxes.

**Extended Data Fig. 3.**
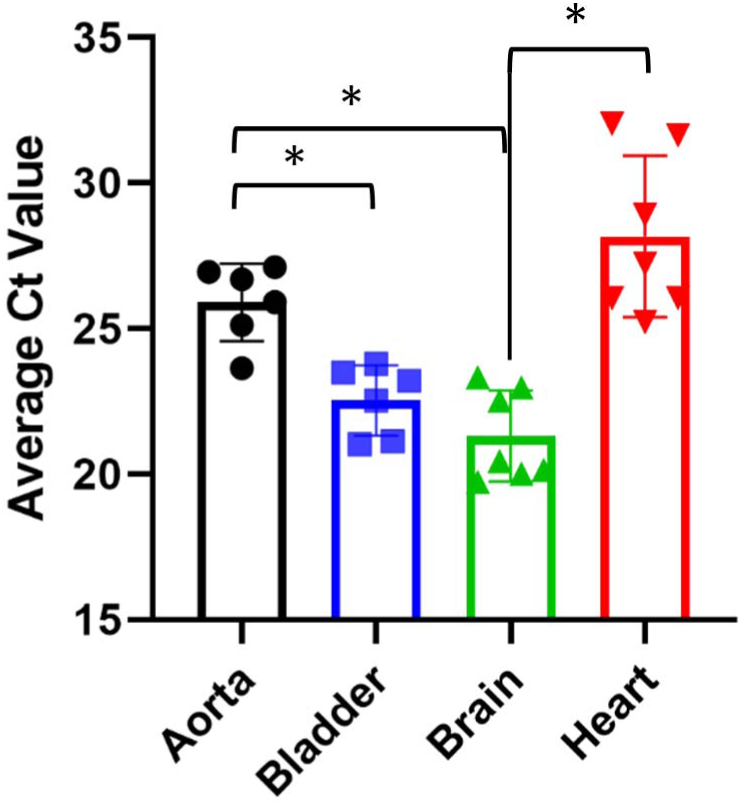
Relative levels of baseline *Tspan2* in mouse tissues. The average Ct value for *Tspan2* mRNA expression is shown in the indicated tissues. A higher Ct value denotes lower expression of *Tspan2*. Data are representative of two independent studies in wild type mice (n= 4 to 6 mice per tissue). The asterisk indicates statistical significance (p < 0.05) between pairwise comparisons following one-way ANOVA and Tukey’s post-hoc test.

**Extended Data Fig. 4.**
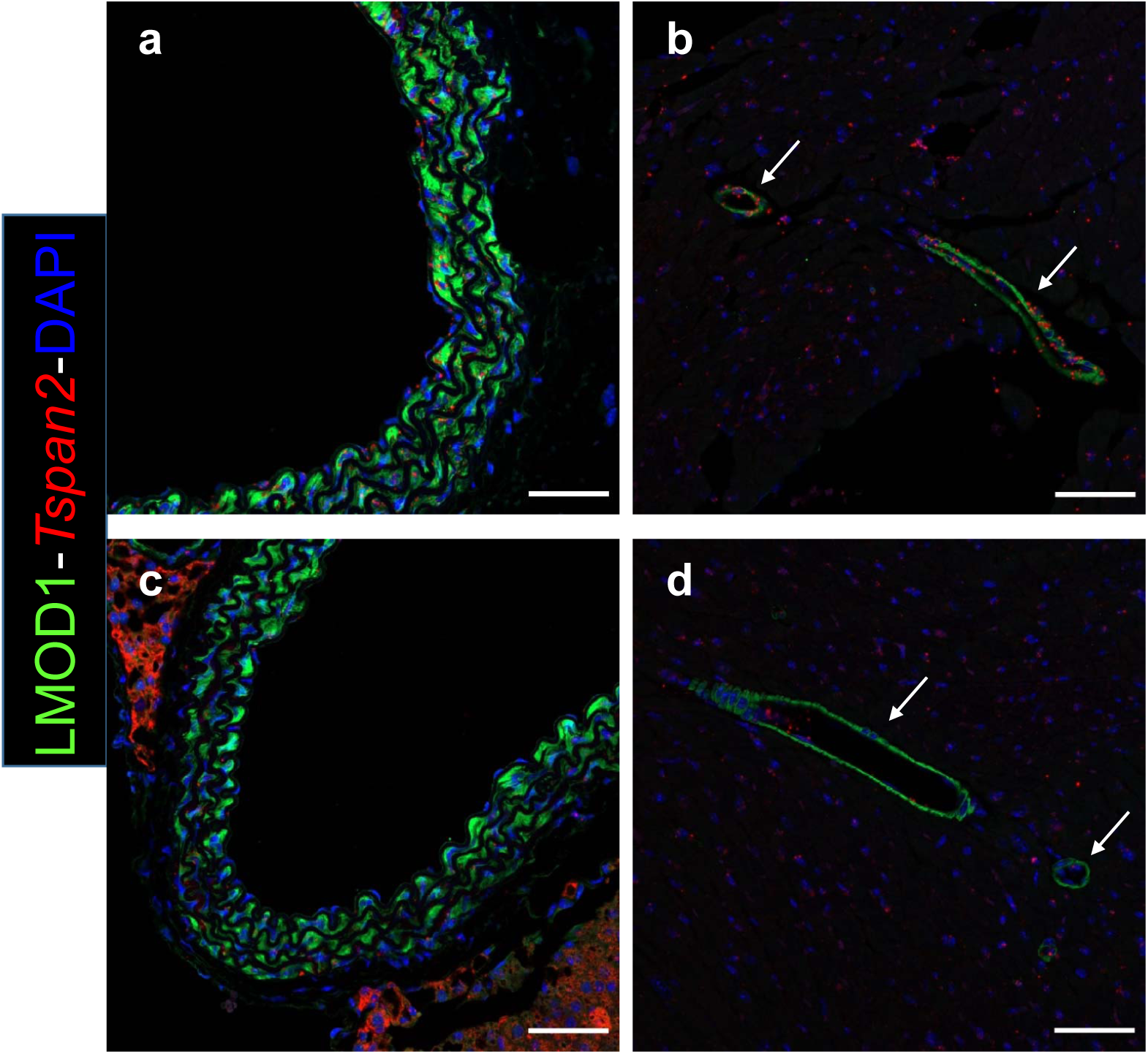
Localization of *Tspan2* mRNA in tissues. Immunofluorescence (LMOD1) and RNA FISH (*Tspan2*) in sections of wild type aorta (**a**) and heart (**b**) versus homozygous *Tspan2* CArG box mutant aorta (**c**) and heart (**d**). Arrows point to coronary vessels of the heart. Note decrease in *Tspan2* mRNA (red dots) in vascular smooth muscle cells (labeled green with LMOD1 antibody) of aorta and coronary vessels in CArG box mutants (**c, d**),

**Extended Data Fig. 5.**
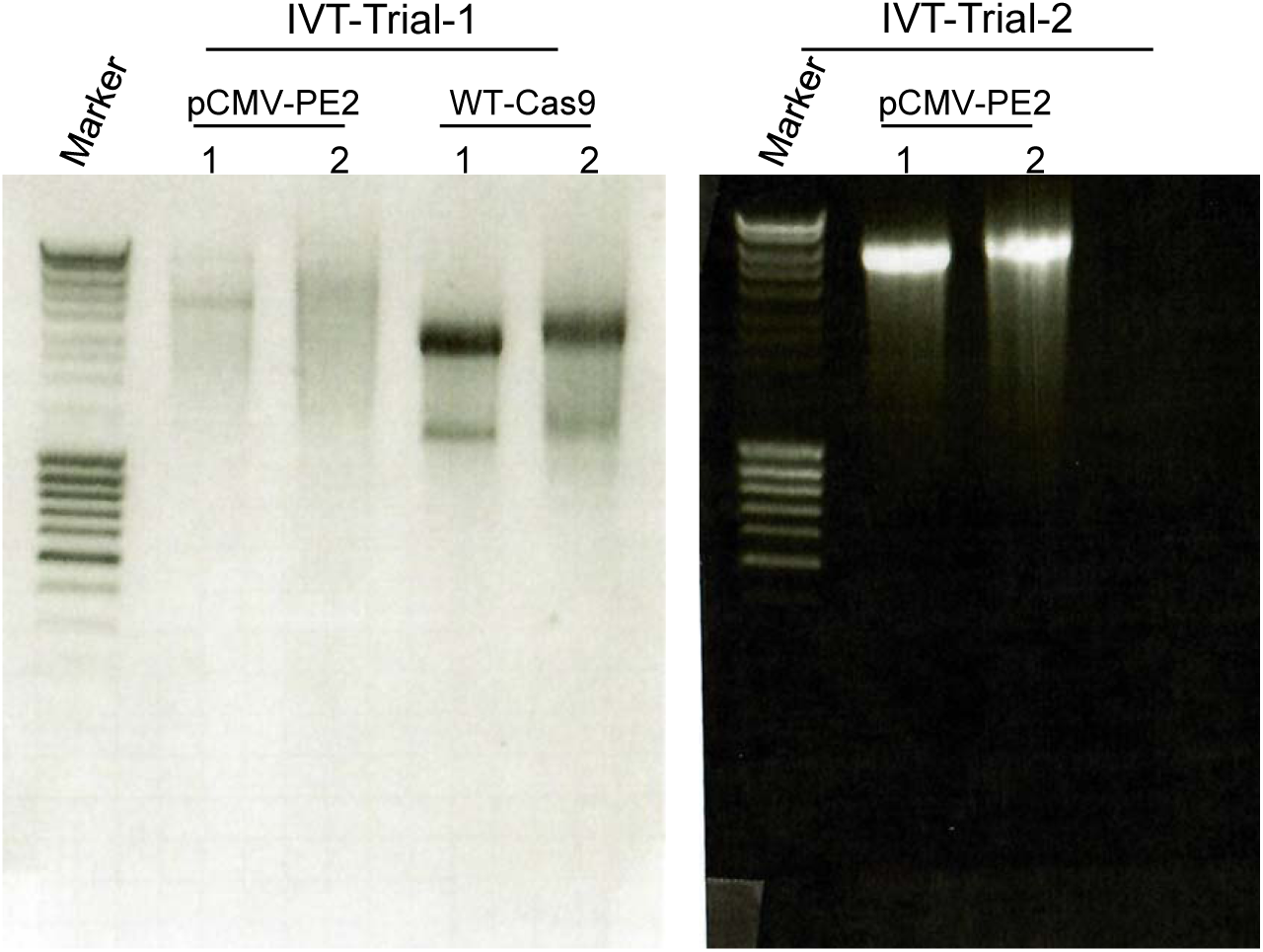
Optimized in vitro transcription of PE2 plasmid. The pCMV-PE2 plasmid was initially In vitro transcribed (IVT) alongside wild type Cas9 with standard conditions per manufacturer (Trial 1). Poor quality PE2 mRNA necessitated a prolonged IVT reaction (3 hrs) and the addition of RNAse inhibitors (Trial 2). The latter conditions resulted in higher quality mRNA for microinjections. Lane 1 samples were untailed and lane 2 samples were polyadenylated.

**Extended Data Fig. 6.**
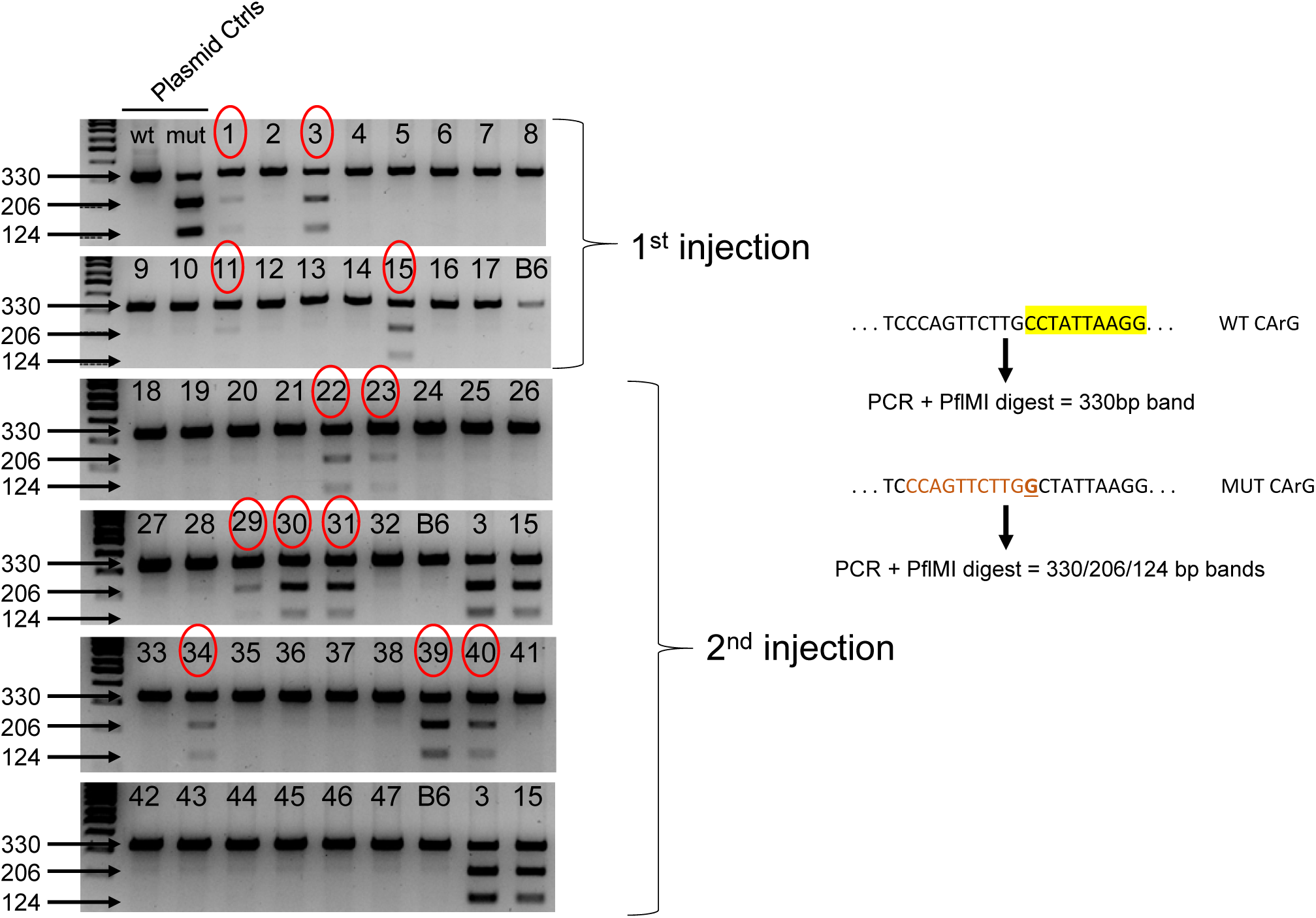
Genotyping of original founder mice derived from PE2 editing of *Tspan2* CArG box. A 330 base pair PCR amplimer was subjected to Van91l (PflMI) restriction digestion (right). The recognition sequence for this enzyme (CCA[N]_5_TGG) is generated with a C>G transversion (bold underlined MUT CArG). Founders circled in red denote those mice used for further analysis. A B6 wild type mouse was used as a negative control and founders 3 and 15 from injection 1 were used as positive controls in founder genotyping of the second microinjection. The CArG box is shown at top right in shaded yellow.

